# Are protected areas tracking threats to terrestrial biodiversity?

**DOI:** 10.1101/2024.10.15.618162

**Authors:** Katherine Pulido Chadid, Carsten Rahbek, Jonas Geldmann

## Abstract

Protected areas (PAs) are vital for nature conservation, yet evidence shows that pressure on biodiversity is increasing despite their global expansion. Using threat probability maps based on the IUCN Red List and PA data, we analyzed the relationship between PA coverage and the major threats—agriculture, hunting, logging, pollution, invasive species, and urbanization—affecting amphibians, birds, mammals, and reptiles. Our analysis includes data on 33,379 species and 255,848 protected sites. Our results reveal a potential disconnect between global PAs and threat tracking, often leaving high-threat areas insufficiently protected. Over half of the mapped area for amphibians and mammals faces high threat impact probability and insufficient PA cover. Amphibians face the highest proportion of high-simultaneous threats and lack sufficient cover. Areas facing a high probability of impact lacking sufficient PA cover also harbor the highest proportion of threatened species across all taxonomic groups. Our research provides crucial insights into the current state of terrestrial PAs concerning threats, highlighting areas requiring immediate attention and guiding strategic conservation planning.

## 2 Introduction

Our planet is experiencing an unprecedented biodiversity crisis, with an increasing number of species facing extinction as we surpass the safe operating boundaries for biosphere integrity (Ceballos et al., 2015; Richardson et al., 2023). The primary threats to biodiversity: land-use change, resource extraction, pollution, invasive species, and climate change, have collectively led to substantial alterations in 75% of Earth’s terrestrial surface (IPBES, 2019). Consequently, at least one-third of vertebrate species have experienced population declines and range contractions (Ceballos et al., 2017), and on average, monitored wildlife populations have experienced declines in the relative abundance of 69% between 1970 and 2018 (WWF, 2022).

In response to this crisis, protected areas (PAs) have emerged as a critical tool for conserving nature in the face of escalating biodiversity decline (Rodrigues & Cazalis, 2020; Watson et al., 2014). Effective PAs can significantly reduce human pressures on ecosystems and safeguard biodiversity (Geldmann et al., 2013; Gill et al., 2024; Wauchope et al., 2022). Recognizing the importance of PAs, international agreements such as the Convention on Biological Diversity (CBD) have continuously called for increased coverage and improved effectiveness of these areas. Although the CBDs strategic plan 2011-2020’s Aichi Biodiversity Target 11 of 17% of protected land by 2020 was not fully met (CBD, 2020), there has been a notable global increase in the coverage of PAs. As of 2024, ca. 17% of terrestrial ecosystems were covered by PAs or Other effective area-based conservation measures (OECMs) (UNEP-WCMC, 2024). With the CBD’s new Kunming-Montreal Global Biodiversity Framework’s (GBF) Target 3, efforts are underway to expand protection further, aiming to cover 30% of the land by 2030 (CBD, 2022).

The ecological effectiveness of PAs depends on their capacity to represent and sustain the diverse ecological features and processes they are designed to protect (Durán et al., 2022; Geldmann et al., 2013). Overall, the extent to which PAs can effectively conserve nature over the long term is determined by how well they deliver positive biodiversity outcomes and abate threats (Geldmann, 2023; Margules & Pressey, 2000; Rodrigues & Cazalis, 2020; Schulze et al., 2018). Several interlinked factors influence this effectiveness: location, spatial design, management, governance, connectivity, and size. Ideally, these factors should lead to the mitigation of threats and enhanced resilience to ensure the long-term persistence of nature within PAs (Durán et al., 2022; Rodrigues & Cazalis, 2020).

Achieving optimal PA effectiveness has proven challenging. Studies indicate limited success in reducing biodiversity loss within PAs, further exacerbated by rising human pressures inside and outside their boundaries, especially in tropical regions (Geldmann et al., 2013, 2019; Jones et al., 2018). The effect of PAs as a tool can also be compromised by placing them in areas with minimal immediate threats. For instance, many PAs have been established in locations with low human pressure - far from roads, cities, and areas of high elevation and low agricultural potential (Joppa & Pfaff, 2009). While these areas might hold significant biodiversity value, they are often under less threat. Thus, the designation of PAs in such areas risks distracting attention away from places more in need of protection and potentially inflating the perceived effect of protection. This is because remote, well-preserved regions with PAs already benefit from inherited protection due to their isolation and distance from threatening processes (Forero-Medina & Joppa, 2010; Joppa & Pfaff, 2009, 2011; Margules & Pressey, 2000).

Most evaluations of PA effectiveness have often concentrated on representation metrics, including the PA network’s overall size, management resource levels, and locational considerations (Geldmann et al., 2015; Rodrigues & Cazalis, 2020). However, recent conservation discussions underscore the importance of prioritizing areas characterized by high threats, inadequate protection levels, and the presence of threatened and/or endemic species rather than only maximizing representation within PAs. By focusing on these areas, conservation efforts can maximize their impact on biodiversity outcomes (Forero-Medina & Joppa, 2010; Joppa & Pfaff, 2009; Negret et al., 2024).

Over the past decade, there has been increasing recognition of threats to biodiversity, leading to the widespread use of threat maps (Tulloch et al., 2015). These maps depict the distribution, intensity, or frequency of threats across landscapes and have guided conservation prioritization by identifying areas where biodiversity is at risk (Brooks et al., 2006; Tulloch et al., 2015). To refine conservation action, threat maps have ranged from evaluating the extent and severity of a single threat to assessing the cumulative impacts of multiple threats on species, using compound indices such as the Human Footprint (Allan et al., 2019; Ostwald et al., 2021; Sanderson et al., 2002; Venter et al., 2016). However, despite advancements, our understanding of context-specific impacts of threats remains limited, either due to insufficient data on specific pressures or the limitations of existing data. For instance, assessments of land-use change derived from remote sensing often reflect physical changes in the environment rather than the ecological consequences for species and ecosystems (Harfoot et al., 2021).

The International Union for Conservation of Nature (IUCN) Red List of Threatened Species is an essential source for assessing the conservation status of species. Through the Red List process, the extinction risk of over 150,000 species of animals, fungi, and plants has been assessed, including the threats affecting each species (IUCN, 2024; Salafsky et al., 2008). However, the Red List lacks information on where these threats occur within a species’ range. The method developed by Harfoot et al. (2021) addresses this challenge by calculating the probability of a random species being affected by a given threat within its distribution while accounting for the uncertainty related to species not necessarily being affected by a threat in all of their range.

While the critical role of PAs in nature conservation is widely acknowledged, efforts to expand their coverage and effectiveness require a deeper understanding of how PAs address threats to biodiversity. Here, we aim to bridge this knowledge gap by analyzing the distribution of the current terrestrial protected cover to the main threats faced by amphibians, birds, mammals, and reptiles across different geographical regions. We do this by conducting a spatial analysis using data from the global network of PAs from the World Database of Protected Areas (WDPA) and threat probability maps for terrestrial vertebrates based on the IUCN Red List (Farooq et al., 2024; Harfoot et al., 2021). We first examine the patterns between PAs coverage and the main threats to terrestrial vertebrates: agriculture, hunting, logging, invasive species, pollution, and urbanization. Based on this, we identify geographical regions across threats and taxa with a high probability of threat and low PA coverage. Then, we examine regions characterized by multiple high-probability threats and insufficient protection. Lastly, we identify the regions of high conservation relevance according to the presence of threatened species. Through this global analysis, we identify patterns in the current state of PA coverage in addressing threats, which will offer valuable insights for future conservation strategies and the ongoing expansion of PAs.

## 3 Methods

### 3.1 Data and preprocessing

#### Threat maps

We employed the methodology established by Harfoot et al. (2021) to develop maps of the probability of impact from the main threats to terrestrial vertebrates. Specifically, we used updated data from the IUCN Red List version 2022-1 and bird species distribution maps of the world—version 2021.1. This data comprises 33,379 terrestrial species, including 7,073 amphibians, 10,959 birds, 5,581 mammals, and 9,766 reptiles (Farooq et al., 2024).

We developed maps for six threats: agriculture, hunting and trapping, invasive and other problematic species, genes & and diseases (From now on referred to as “invasive species”), logging, pollution, and urbanization. This yielded 24 maps, each corresponding to the probability of threat for a specific taxonomic group and threat. All maps had a 50 × 50 km resolution using an equal-area Mollweide projection. Following Harfoot et al. (2021), only cells with species richness greater than ten were considered in the analysis. The dataset was then linked with its corresponding Biogeographical realms (ArcGIS Hub, 2009) and United Nations (UN) geographical regions and subregions (United Nations, 2024). In this process, Oceania was merged into Australasia, and the Antarctic was removed due to limited information on threat data.

In total, our maps covered 54,685 cells for birds (∼ 136.71 million km2 of the land surface), 50,624 for mammals (∼ 126.56 million km2), 36,252 for reptiles (∼ 90.63 million km2) and 22,936 for amphibians (∼ 57.34 million km2). The number of cells containing data per taxa differed, reflecting the data availability and ranges of the species studied.

#### Terrestrial protected areas

The WDPA is the most comprehensive global repository of PAs and OECMs compiled through collaborative efforts of governments and stakeholders. We obtained data on terrestrial PAs from WDPA, sourced from the September 2023 release. This dataset contains polygon information for 273,898 terrestrial, coastal, and marine sites (UNEP-WCMC and IUCN, 2023).

As preprocessing steps, we first filtered the data to retain PAs within terrestrial and coastal regions. We then intersected the remaining sites with land surface data from (OpenStreetMap, 2019). Sites meeting the definitions of PAs or OECMs, categorized as designated (legally/formally designated), established (established through other effective means) and not reported, were selected for analysis (UNEP-WCMC, 2019). Due to restricted access to data on PAs, China, India, and Turkey were not available in the WDPA. Thus, these countries were excluded from the analysis. After these steps, our final dataset included 255,848 sites.

To calculate the percentage of a cell covered by PAs, the polygons were rasterized into 50 × 50 km grid cells, matching the projection, resolution, and extent of the threat probability maps. The coverage of each cell was then determined as the proportion covered by PAs within each cell. The spatial operations were developed using R version 4.2.3 (R Core Team, 2021) and the R packages sf (Pebesma, 2018), raster (Hijmans, 2023a), and terra (Hijmans, 2023b).

### 3.2 Data analysis

#### Assessing the overall global patterns between PA coverage and threat probabilities

To look at the global patterns of PA coverage and the probability of different threats, we combined the layer of PA coverage with layers of threat probabilities. Our initial data exploration revealed a non-linear relationship between PA coverage and threat probability across taxa. Due to the high number of observations, linear regression might be susceptible to overplotting, making visualization and interpretation of trends difficult (Mayorga & Gleicher, 2013). To address this challenge, we employed a Generalized Additive Model (GAM) approach to illustrate the relationship between protection and threat probabilities. GAMs are well-suited for analyzing non-linear relationships between variables. Unlike linear models that assume a straight-line relationship, GAMs allow for the inclusion of smooth functions to capture complex, curved patterns in the data (Hastie & Tibshirani, 1986).

#### Exploring the influence of protected area age on threat levels

We assessed the relationship between the year of designation and the maximum threat value for each site and taxa to examine any relationship between how long a site had been protected and the probability of threats. We used the median threat value across each site per taxa and tested the correlation between these variables. As a data-cleaning step, sites without the year of establishment and cells without threat values were excluded from the analysis.

We found no strong global trend. Using Spearman coefficient, we observed weak positive correlations between PA designation year and maximum threat level for reptiles (rho = 0.16), amphibians (rho = 0.082), and birds (rho = 0.061). This suggests a slight tendency for Pas established later to have higher threat levels for these taxa. For mammals, a weak negative correlation was found (rho = -0.056), suggesting a possible slight decrease in the maximum threat levels over time.

#### Analyzing the proportions of threats and protected areas across the globe

We analyzed the distribution of threats relative to PA coverage. We categorized threat probability and PA coverage into consistent discrete groups to facilitate this analysis and ensure comparability across different groups (Table S1). While the specific thresholds chosen for threat categories involve some level of discretion, we defined cells under a high probability of impact (“high threat”) as those where the threat could impact at least 25% of the species in a given cell.

The PA coverage categories were chosen to reflect past and current policy objectives to address global biodiversity loss. “Low” coverage refers to areas falling below the 2020 Aichi Target of 17% protected land, a goal established by the CBD in 2010. “High” coverage, on the other hand, aligns with the Kunming-Montreal GBF target of 30% protection by 2030. We also use the term “insufficient cover” to refer to cells with unprotected and low-protected cover.

To determine the extent of land that faces various levels of threat and the degree of protection it receives, we analyzed the distribution of different categories and computed the percentage of each category for each threat and taxa. We also calculated the average percentage across all taxa.

#### Critical geographical regions within and across taxa

We analyzed the spatial distribution of threats and PA cover categories to identify critical regions for conservation within and across taxa. We aimed to detect the areas with the highest probability of threat impact. For this, we extracted grid cells where at least one threat category was classified as “high”, thus extracting the maximum threat value per grid cell. Additionally, we analyzed the regional patterns with a high probability of impact and insufficiently protected cover within and across taxa.

To further identify areas in potential need of enhanced protection, we calculated the frequency of high-threat occurrences within unprotected cells for each taxon. Then, focusing solely on areas facing four or more simultaneous threats per taxon, we identified sub-regions where at least 5% of the total area experiences multiple threats across different species groups.

Finally, to identify areas potentially in urgent need of conservation efforts, we examined the proportion of threatened species across all threat and PA coverage categories. Thus, we calculated the proportion of species categorized as Vulnerable, Endangered, or Critically Endangered (IUCN criteria) relative to the total number of species in that cell. This analysis identified areas where the proportion of threatened species exceeded a pre-defined threshold of 15%.

### 3.3 Sources of uncertainty

The method employed by Harfoot et al. (2021) estimates threat probability across a species’ entire range. However, this approach can introduce uncertainty, particularly for species with large ranges. Since the method assumes threats are present throughout the range, reliability decreases as the range size grows. We calculated the median range size within cells and taxa to identify areas where species have higher median ranges and, thus, where potential uncertainty could be higher.

Additionally, species richness is generally higher in the tropics than in temperate regions, contributing to potential sources of uncertainty when the probability of impact is calculated based on the number of species within a grid cell. Therefore, areas with low species richness could overestimate threat impact due to the averaging effect across fewer species. To address this, we identified areas with the largest median range sizes and the lowest species richness.

## 4 Results

### The global patterns between protected area coverage and threats

Agriculture, hunting, logging, and urbanization were the major threats across all four taxa, with amphibians being the most affected. However, the importance of these four threats varied across taxa. Agriculture was among the top three threats for all taxa, particularly affecting amphibians and reptiles. Likewise, hunting was among the top two threats for all taxa except amphibians. Logging ranked second for amphibians and third for mammals, and urbanization was the top threat for both amphibians and reptiles (Figure 1).

**Figure 1.**
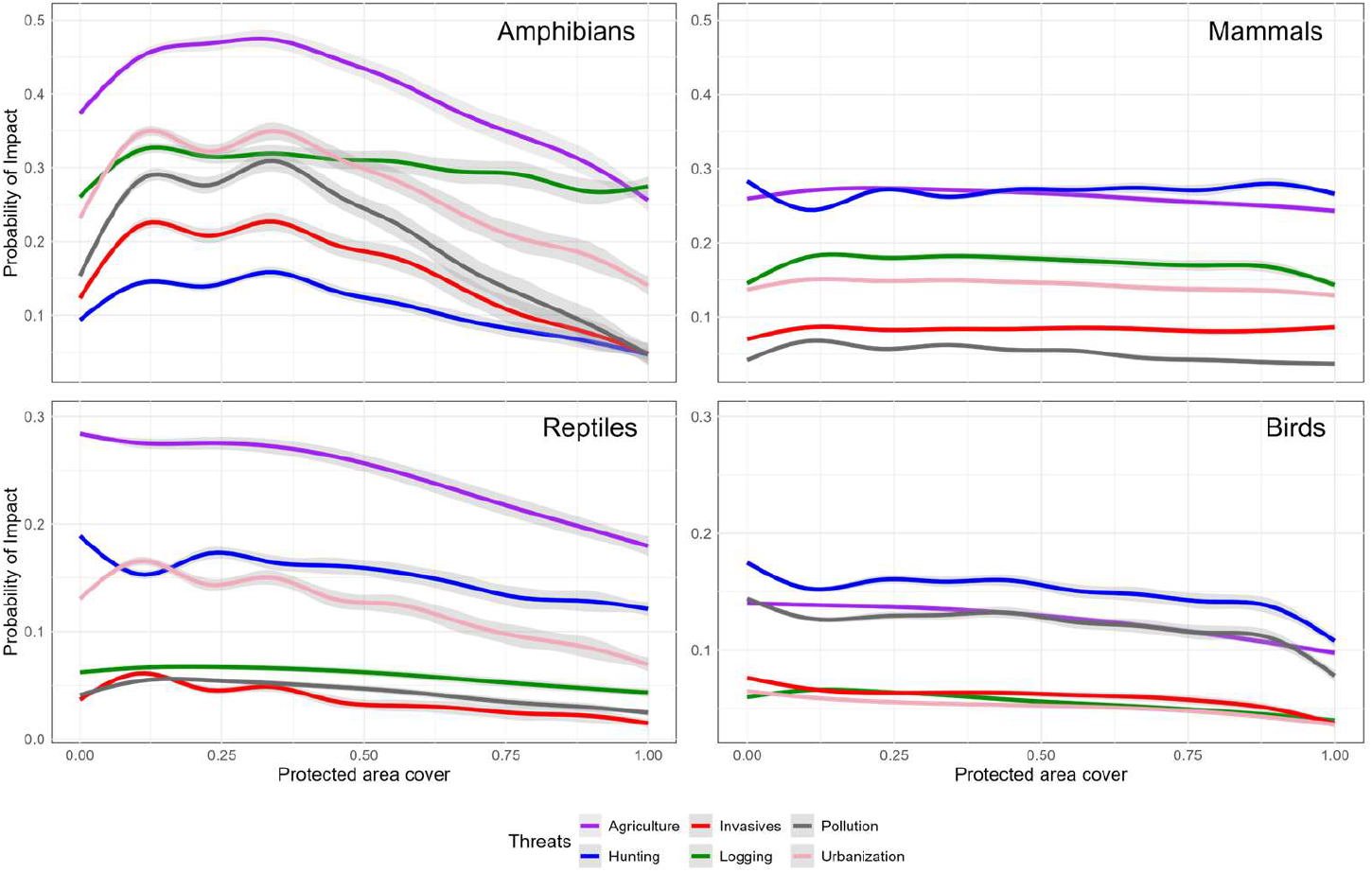
Relationship between protected area coverage and probability of impact per threats and taxon

When looking at the relationship between the probability of threat impact and the percentage of protection, there was, in most cases, no strong relationship, particularly for mammals and birds. For amphibians and reptiles, however, the impact probability exhibited an inverse relationship with PA coverage, so that areas with the highest levels of protection had the lowest probabilities of impact from any of the six threats (Figure 1).

### Land facing varying threat probabilities and their protected area allocation

Areas facing the highest probability of impact were mostly unprotected or had low protection (Figure 2). However, across all levels of threat, protected cover remains relatively constant, with 75%, 77%, and 76% of areas having low or no protection for low, medium, and high threat levels, respectively (Figure 2). Cells with high-impact probability covered 21% of the mapped terrestrial surface of Earth, but only 16% of this area (2.97 million km²) had more than 30% PA coverage, while areas with lower threat probability had a slightly higher proportion of highly protected land (18.5% with >30% PA coverage; Figure 2).

**Figure 2.**
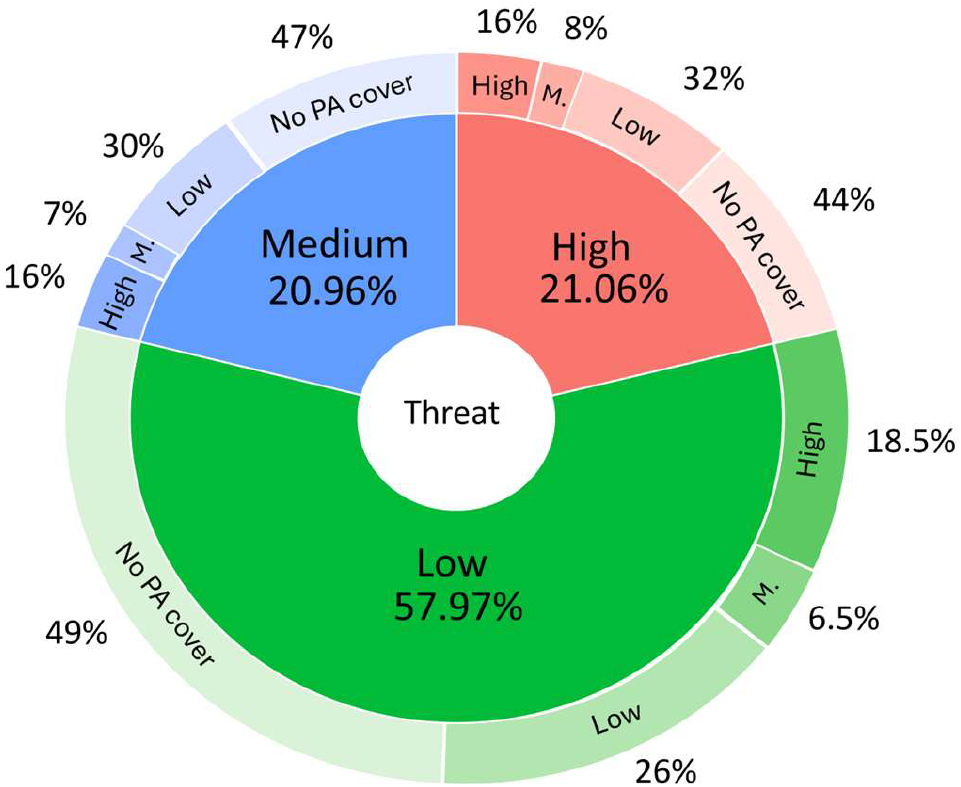
Average probabilities of the total low, medium, and high threat probability across all threats and levels of PA coverage (No cover = 0; Low protected cover = < 17%; Medium cover = 17 – 30%; High protected cover = 30%).

### Geographic patterns of threats and protected cover

Globally, 58.7% of cells for mammals, 54.6% of cells for amphibians, 40.7% of cells for reptiles, and 22.3% of cells for birds had at least one threat with a high probability of impact in areas of low or absent protection (Figure 3). The geographic distribution of these threats varied across taxa and threat types. Overall, threats to amphibians, agriculture, and hunting for birds, mammals, and reptiles had a broad geographic extent across all taxa. In contrast, invasive species, particularly birds and reptiles, are mostly restricted to specific regions dominated by islands such as Micronesia and Polynesia (see Supplementary Figures S1 – S4 for details).

**Figure 3.**
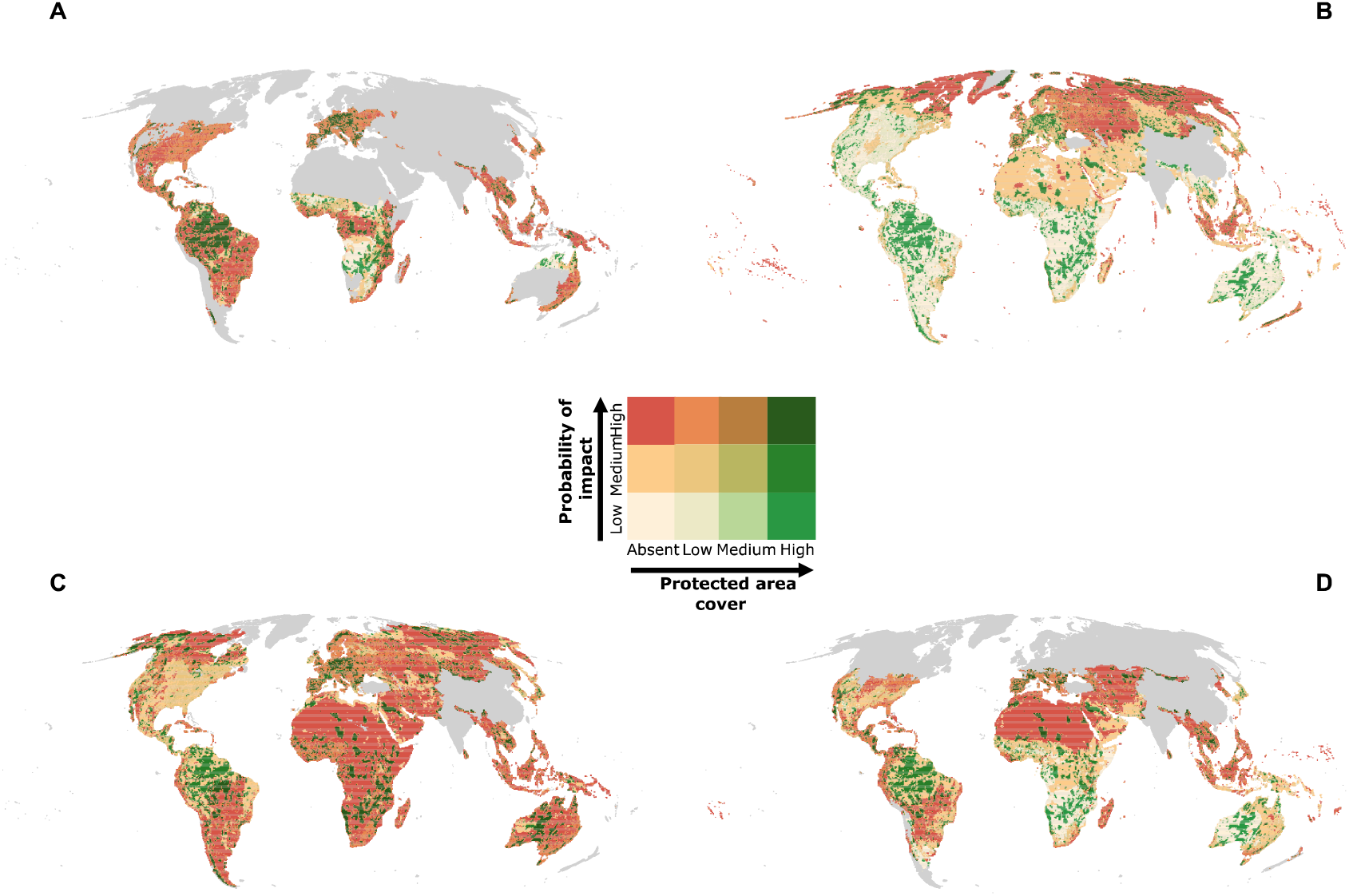
Maximum probability of impact of any threat per given cell for (A) Amphibians, (B) Birds, (C) Mammals, and (D) Reptiles. Threat level and PA coverage are represented by color gradient, with darker tones representing higher levels.

Our results show that over 43% of the mapped area for amphibians, 32% for mammals, and 25% for reptiles faced more than two simultaneous high threats in insufficiently protected cells. However, findings for amphibians are even more concerning, with nearly 16% of their mapped area exhibiting overlaps of four or more co-occurring threats in these cells (Figure 4). The spatial patterns of these areas were highly variable, with no single sub-region emerging with more than four threats for all taxa. Amphibians were the taxa with the highest number of subregions impacted by over four threats (Figure 4, E).

**Figure 4.**
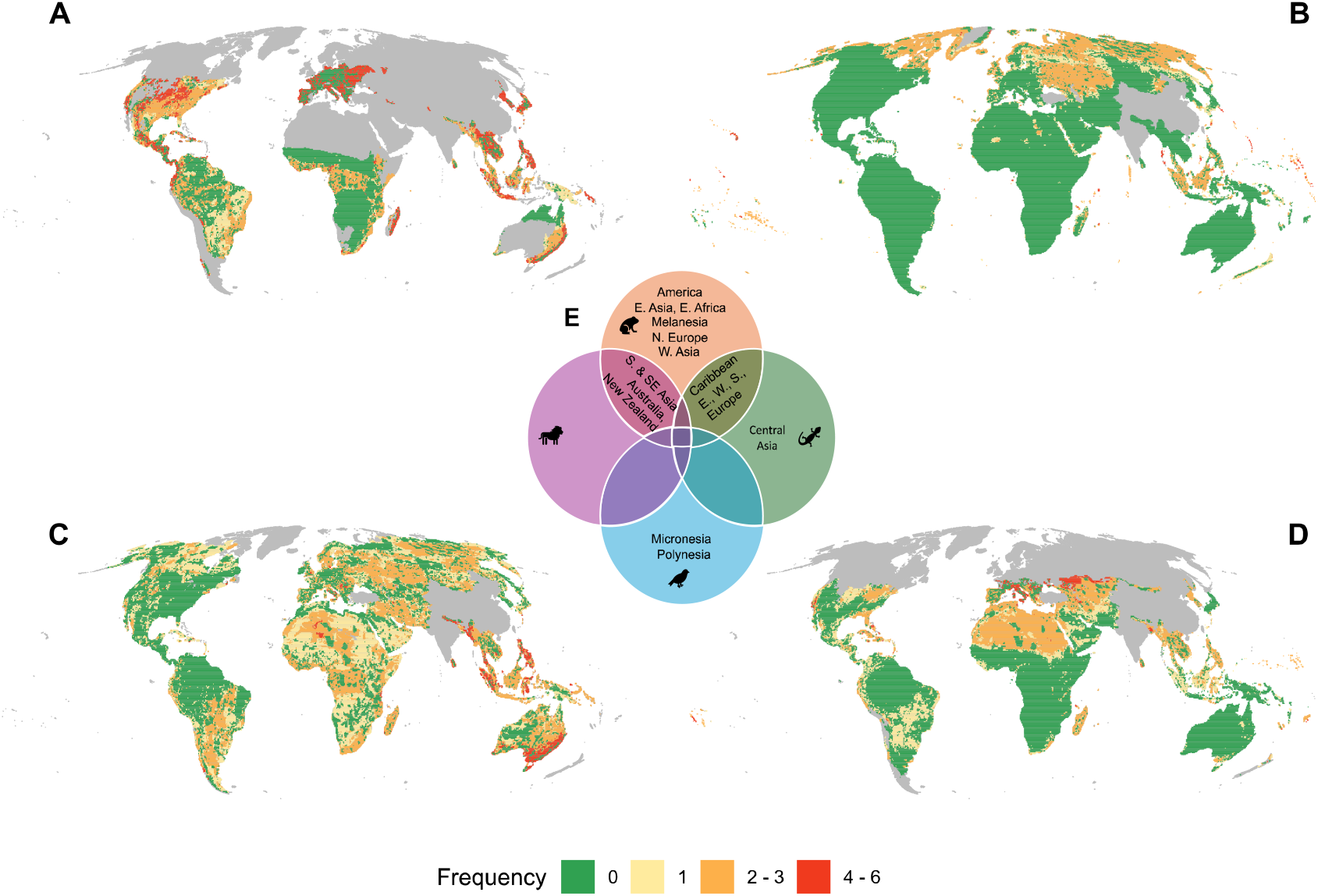
Frequency of areas facing from none to multiple high-impact probability threats and insufficient PA coverage (A. Amphibians, B. Birds, C. Mammals, D. Reptiles, E. Subregions where at least 5% of the total area faces over four simultaneous high-threats and have insufficient protection). Grey cells represent areas with no data

### Potential areas of conservation concern: High threat levels, high proportions of threatened species, and low protected cover

Cells containing more than 15% of threatened species were primarily located within insufficiently covered areas with the highest probability of impact. These areas were particularly found in Central America and the South American Andes for amphibians, the Caribbean for both amphibians and reptiles, Polynesia and Micronesia for birds, and Madagascar and Southeast Asia for mammals (Figure 5).

**Figure 5.**
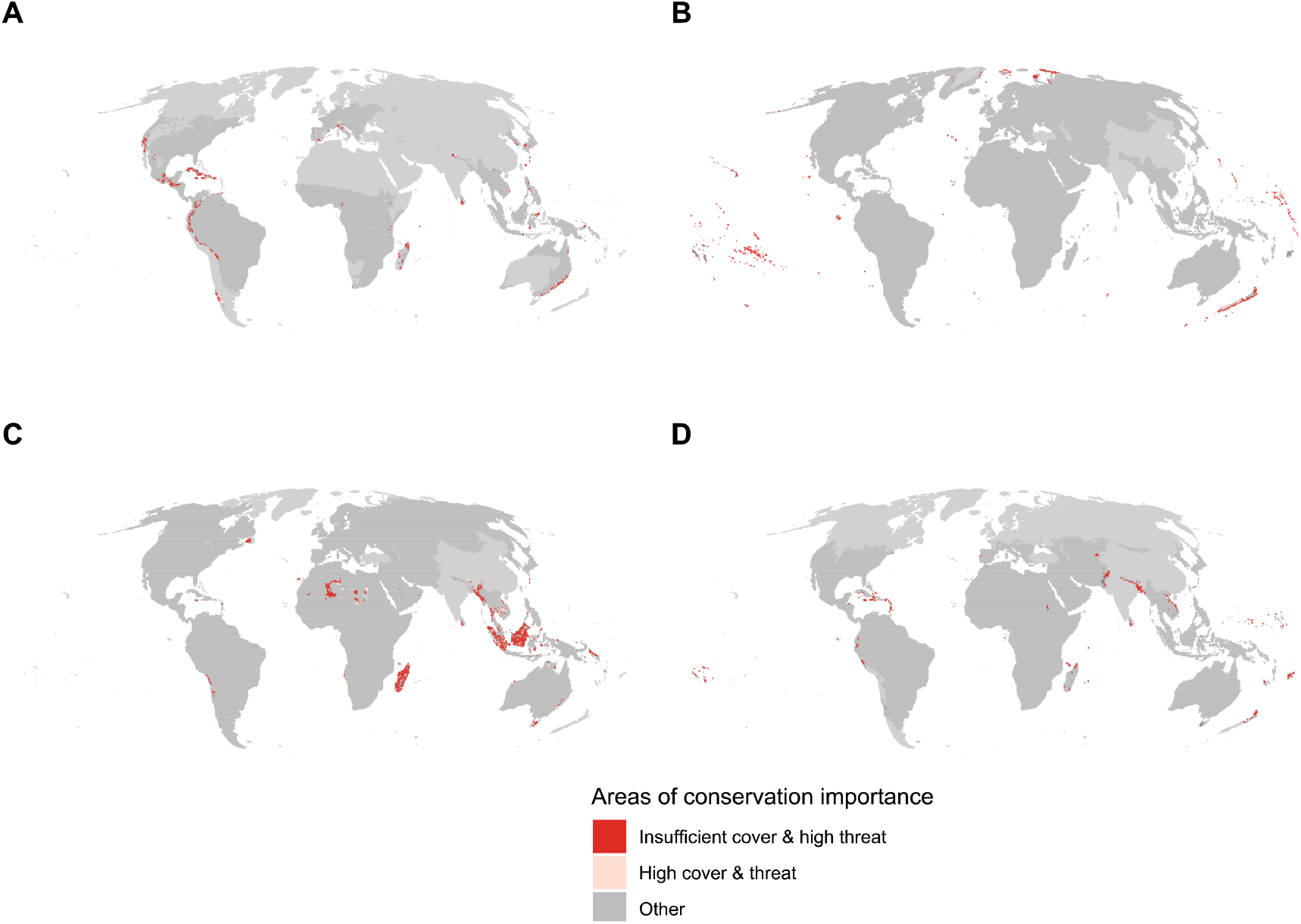
Areas of conservation relevance (Red areas correspond to cells where: 1. Protected area coverage is low or absent (<17%), 2. There is a high probability of threat impact, and 3. The proportion of threatened species is higher than 20% relative to the species richness of the cell. (A. Amphibians, B. Birds, C. Mammals, D. Reptiles)

## 5 Discussion

### Global protected areas are not tracking threats to terrestrial vertebrates

Our global analysis using data from 33,379 species and 255,848 PAs revealed that PAs are not tracking the main threats faced by terrestrial vertebrates. Consequently, most of the land surface facing the highest probability of impact was insufficiently protected. Our results support earlier findings that conservation efforts often prioritize establishing PAs in less productive or remote locations, potentially neglecting areas where wildlife needs the most protection (Joppa & Pfaff, 2009; Vieira et al., 2019).

One reason for this could be that many conservation strategies prioritize regions with high irreplaceability and low threats where conservation interventions are more likely to appear successful, while less emphasis has been placed on vulnerability, and thus, in regions facing the highest threats (Brooks et al., 2006; Joppa & Pfaff, 2009). However, sites of high irreplaceability and high threat require immediate conservation attention to prevent the loss of unique biodiversity values, as replacement options are spatially or temporally unavailable (Rodrigues et al., 2004).

### The challenge of diverging threat impacts across taxa

Agriculture, hunting, logging, and urbanization were the top threats identified for terrestrial vertebrates, but their potential impact on species and their extent differ according to each taxonomic group. Previous research has highlighted that habitat loss and degradation are the main factors threatening amphibians and mammals (Ficetola et al., 2015; Schipper et al., 2008) and hunting to mammals (Schipper et al., 2008), patterns that have also been observed inside PA (Schulze et al. 2017). On the other hand, we also found that invasive species have a focalized occurrence in birds and reptiles. Studies have highlighted that invasive species on islands with insufficient protected cover can have devastating local impacts (Blackburn et al., 2004). However, threat ranking provides a global picture of threat allocation, but the specific importance of each threat can vary depending on the local context and the species (Bellard et al., 2022).

### High-probability threat areas remain largely unprotected, particularly for amphibians

Over half of the mapped area of amphibians and mammals overlapped with at least one high-impact threat in places where PA coverage is insufficient. And for amphibians, many areas are even facing multiple high-impact threats. Our results, thus, confirm previous studies showing that amphibians is one of the most threatened and overlooked groups (Beebee & Griffiths, 2005; Stuart et al., 2004a). Previous studies have also shown that amphibians receive notably less conservation attention than other taxa (Creighton & Bennett, 2019; Nori & Loyola, 2015; Rodrigues et al., 2004). This disparity is likely attributed to the generally smaller geographic ranges of amphibians, making them more susceptible to being excluded in PAs, and a taxonomic bias in conservation efforts, with fewer sites specifically designated for amphibian conservation compared to those for birds and mammals (Rodrigues et al., 2004).

We observed that 43%, 32%, and 25% of the mapped area for amphibians, mammals, and reptiles had multiple threats. Out of these, 16% of the mapped area for amphibians experienced four or more simultaneous high threats in areas insufficiently protected. This is concerning, as multiple threats can exacerbate the impact further (Beebee & Griffiths, 2005; Hof et al., 2011; Menéndez-Guerrero & Graham, 2013; Stuart et al., 2004a). Thus, the presence of co-occurring threats can act as a major stressor, intensifying the loss of resilience in vertebrate populations and the loss of species with critical functional traits, which is likely to disrupt ecosystem functions (Bellard et al., 2022; Capdevila et al., 2022; Munstermann et al., 2022).

### Areas with insufficient protection and high threat levels host most threatened species

Our analysis revealed geographic heterogeneity in the distribution of high-threat areas with insufficient protection across taxa, and no single subregion emerged as a critical region for all groups. Alarmingly, these high-threat areas coincide with the highest concentrations of threatened species. The dispersed distribution of areas under high impact probability across taxa is a challenge for conservation planning and may also be one reason why so many areas under high threat remain insufficiently protected. Identifying spatial conservation priorities is a complex process, with numerous options and criteria to be considered for creating or expanding PAs—a process often constrained by limited resources (Critchlow et al., 2021). In addition, current PAs allocation has been based on limited species, leading to the underrepresentation of rare and non-target taxa, which affects conservation outcomes (Delso et al., 2021; Kujala et al., 2018; Westgate et al., 2014).

The aforementioned could explain the differences between protected cover, threats, and taxa found in our analysis. Furthermore, the gap between research and implementation often hinders effective conservation planning (Knight et al., 2008). Thus, the ongoing trend of favoring low-cost lands for protection could also be attributed to a lack of political will to protect high-cost areas (Venter et al., 2018). Urgent action is needed, with increased efforts to reverse these trends. This is particularly important in Central America and the South American Andes for amphibians, the Caribbean for both amphibians and reptiles, Polynesia and Micronesia for birds, and Madagascar and Southeast Asia for mammals. Previous research has highlighted these areas as experiencing high rates of biodiversity loss across taxa (Farooq et al., 2024; Schipper et al., 2008; Stuart et al., 2004b).

### Uncertainty in threat probability maps

While our research offers insights into the threats faced by different taxonomic groups in relation to PA coverage, interpreting the results requires careful consideration due to inherent uncertainties associated with the employed threat probability maps. These maps primarily rely on IUCN Red List species range data, representing potential species distributions (Di Marco et al., 2017; Rocchini et al., 2011). Several factors contribute to uncertainties: limitations of IUCN Red List data due to reliance on inference and expert judgment (IUCN, 2024; Hayward et al., 2015), commission and omission errors in species range maps (Di Marco et al., 2017; Rodrigues et al., 2004; Ficetola et al., 2014), and location bias favoring well-studied regions (Rocchini et al., 2011). Furthermore, uncertainties extend beyond species distribution. The threat probability maps represent the likelihood of where species are likely to be affected by a certain threat, but not necessarily on its severity and species-specific vulnerability (Farooq et al., 2024; Harfoot et al., 2021)

Another key limitation lies in the method’s assumption of uniform threat probability across a species’ entire range. This, coupled with large species ranges and low species richness, can introduce uncertainties into the results (Figures S9 & S10). For instance, the high threat probability for mammals and reptiles due to agriculture and hunting in the Sahara Desert, or the high threat probability for birds due to hunting and pollution in the high Arctic, might be a consequence of species threatened elsewhere in their range contributing to the overall threat score in those locations. The low species richness of these regions could also lead to an overestimation of threat values for the species present. Therefore, these high threat probabilities may not necessarily reflect the presence of highly likely threats within these regions.

### Uncertainty related to protected areas

Our study utilized data on PA coverage to identify spatial patterns of threats and ongoing conservation efforts. However, it is crucial to acknowledge the limitations of our approach. Firstly, the WDPA might contain inaccuracies. Some entries may represent “paper parks” that lack sufficient resources or enforcement (Juffe-Bignoli et al., 2014). These PAs can inflate the perceived effectiveness of PAs, potentially masking areas where threats remain high despite designated protection. In addition, the WDPA relies on national reporting, which can be inconsistent or vary in quality (UNEP-WCMC, 2019). This introduces uncertainty about the actual ecological functionality of PAs within a region. A high coverage area based on national reports might not translate to solid protection on the ground.

While our study primarily examined PA coverage to identify spatial patterns of threats and conservation efforts, it is important to acknowledge that PA coverage alone does not guarantee effectiveness (Geldmann et al., 2019; Ghoddousi et al., 2022; Jones et al., 2018). Factors like funding, staffing, and enforcement play a critical role in a PA’s ability to protect biodiversity (Bruner et al., 2001; Correa et al., 2024; Graham et al., 2021). A small, well-managed PA might provide better protection than a larger PA with inadequate resources. Thus, further research is needed to evaluate the effectiveness of existing PAs in mitigating these identified threats.

We found slight and weak correlations between PA age and threat probabilities. Reflecting that older PAs, does not necessarily relate to lower values probabilities of threat at the global level. This potentially illustrates an additional challenge of separating the effects of PA location from PA performance on threat mitigation (Ferraro et al., 2019; Geldmann, 2024). Our approach may not distinguish between areas with low threats due to effective PA management in high-threat areas or PAs that have been placed in areas with low levels of threats to begin with. However, while it is difficult to separate the causes for the observed patterns, this does not change the fact that too many areas of high threat remain insufficiently protected. Thus, our results can still help to guide future PA strategies.

Lastly, the methodology used in our analysis focuses on the percentage of each cell being protected rather than assessing the actual coverage of PAs within the broader landscape. This distinction is crucial because it implies that our analysis might overlook spatial details beyond our relative coarse resolution of 50×50 km. For instance, the most threatened areas within a particular cell could lie outside the regions designated as protected even if the percentage of that protected cell exceeds 30%. This discrepancy arises because the method aggregates data at the cell level without accounting for the precise locations of protected and unprotected areas within each cell.

#### 5.1 Perspectives

Our global analysis is not intended to guide policy or conservation action in specific countries (Chaplin-Kramer et al., 2021; Joppa & Pfaff, 2011). However, our study builds upon understanding how PAs are currently allocated relative to threats faced by different taxa. This analysis serves two key purposes. First, it gives us an overview of the current conservation efforts across various regions and taxa. Helping to identify areas where immediate action might be necessary to prevent extinctions. Second, it provides a global perspective that can help inform national and local decisions on potential areas where future interventions are critical for achieving global biodiversity goals, such as the 30×30 initiative.

To reverse current extinction trends, increased protection remains crucial for safeguarding biodiversity and enhancing PAs’ effectiveness, especially in regions where species are threatened. The establishment of PAs should, thus, target areas and species likely to be lost without protection (Negret et al., 2024; Pressey et al., 2015). Also, scaled-up investment is needed to improve the effectiveness of existing PAs and address threats in these high-risk regions (Luedtke et al., 2023).

## Notes

### Competing Interest Statement

The authors have declared no competing interest.

