## Supplementary material for "Are protected areas tracking threats to terrestrial biodiversity?"

Table S1. Categories for threat probabilities and PA coverage

| Threats | | Protected areas | |
| --- | --- | --- | --- |
| Probability of impact (%) | **Category** | **Coverage (%)** | **Category** |
| - | - | 0 | Unprotected |
| 0 - 12.5 | Low | < 17 | Low |
| 12.5 - 25 | Medium | 17 - 30 | Medium |
| 25 - 100 | High | 30 - 100 | High |

Table S2. Proportion of total area covered by low and high probability of impact respective to protected coverage (%)

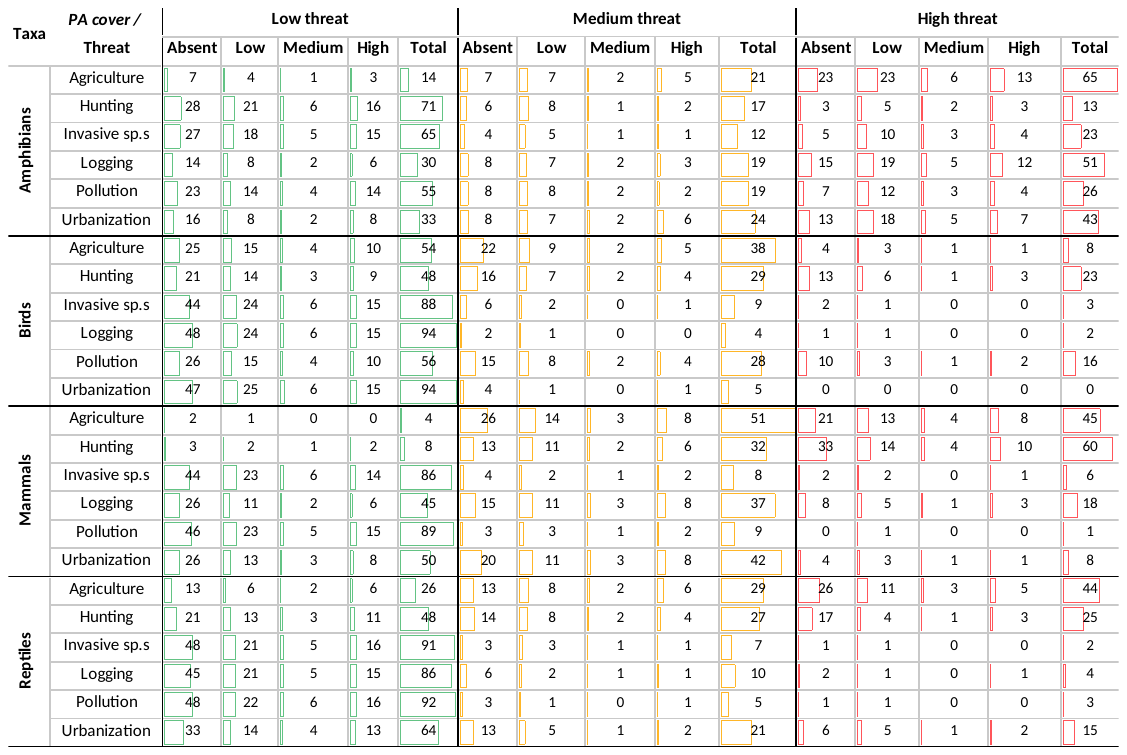

Values indicate the percentage of total area covered by low, medium and high probability of impact respective to protected coverage (%).
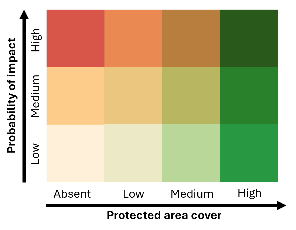

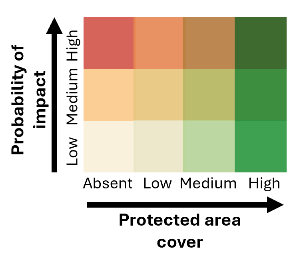

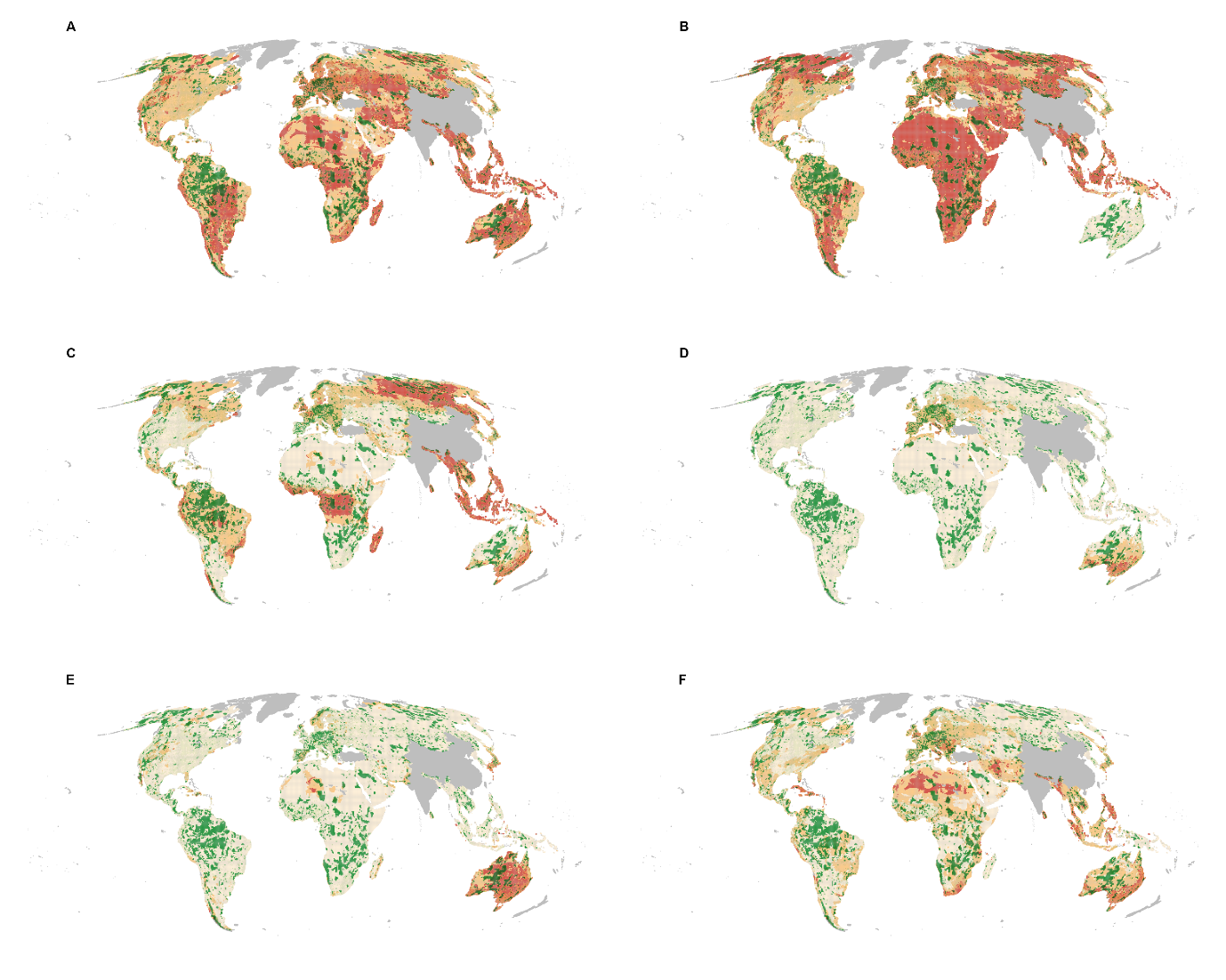

Figure S1. Bivariate maps for PA cover and threats to mammals (A. Agriculture, B. Hunting, C. Logging, D. Pollution, E. Invasive species F. Urbanization)

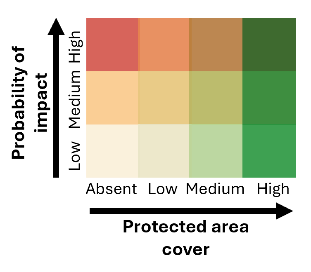

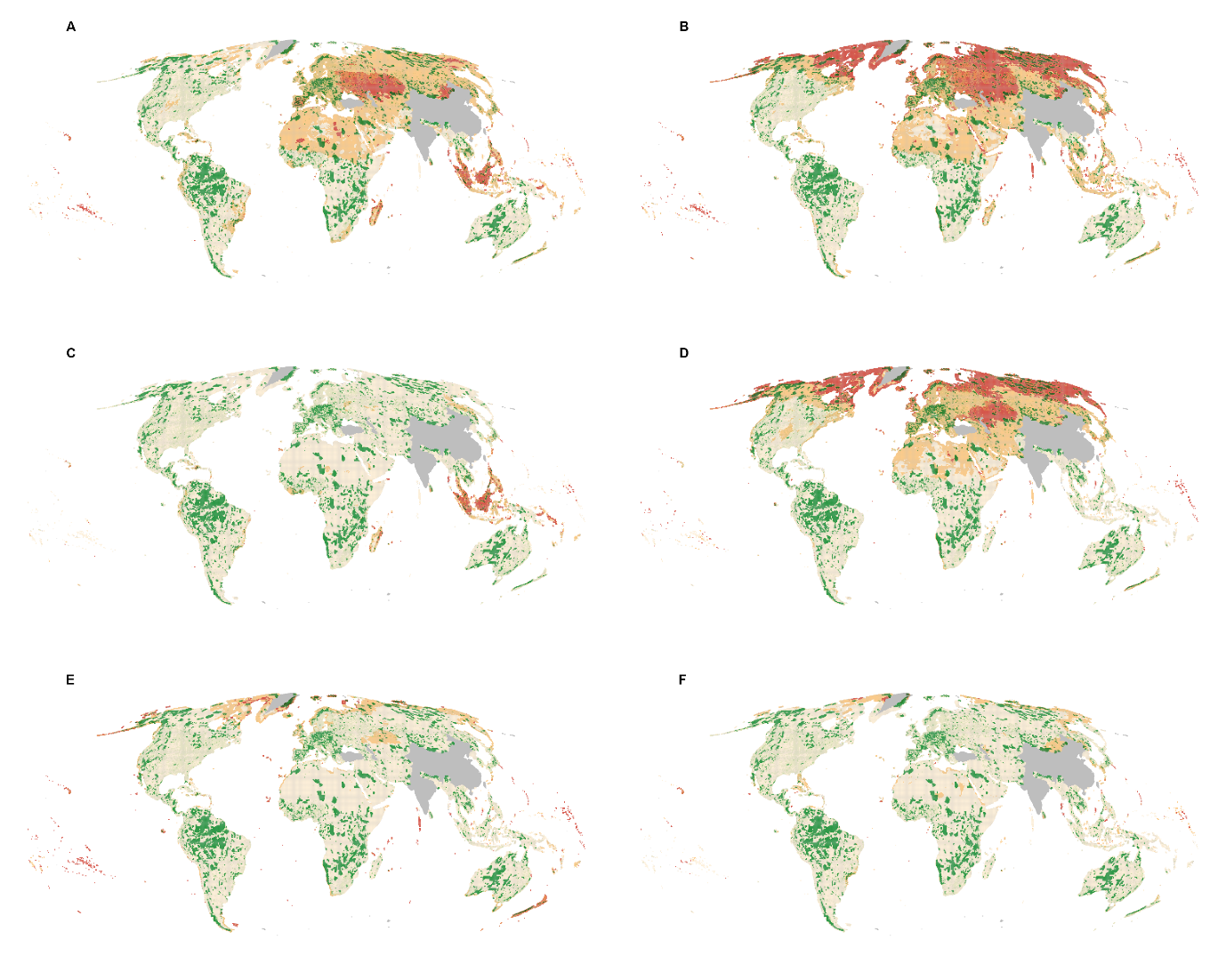

Figure S2. Bivariate maps for PA cover and threats to birds (A. Agriculture, B. Hunting, C. Logging, D.Pollution, E. Invasive species F. Urbanization)

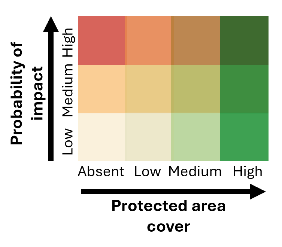

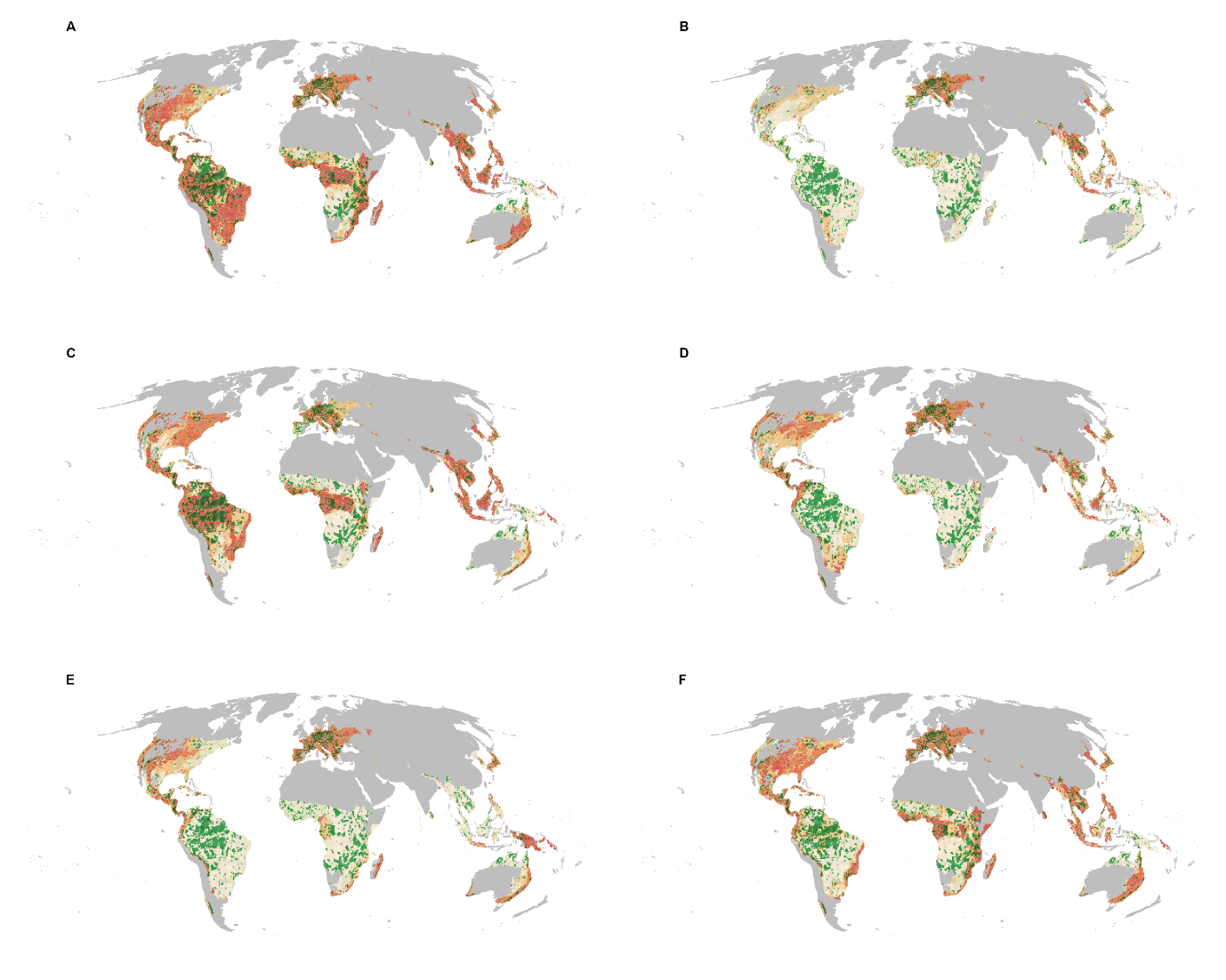

Figure S3. Bivariate maps for PA cover and threats to amphibians (A. Agriculture, B. Hunting, C. Logging, D. Pollution, E. Invasive species F. Urbanization)

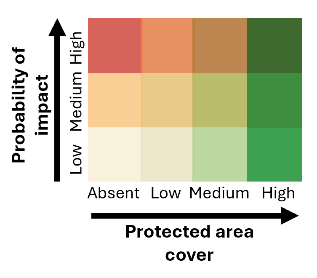

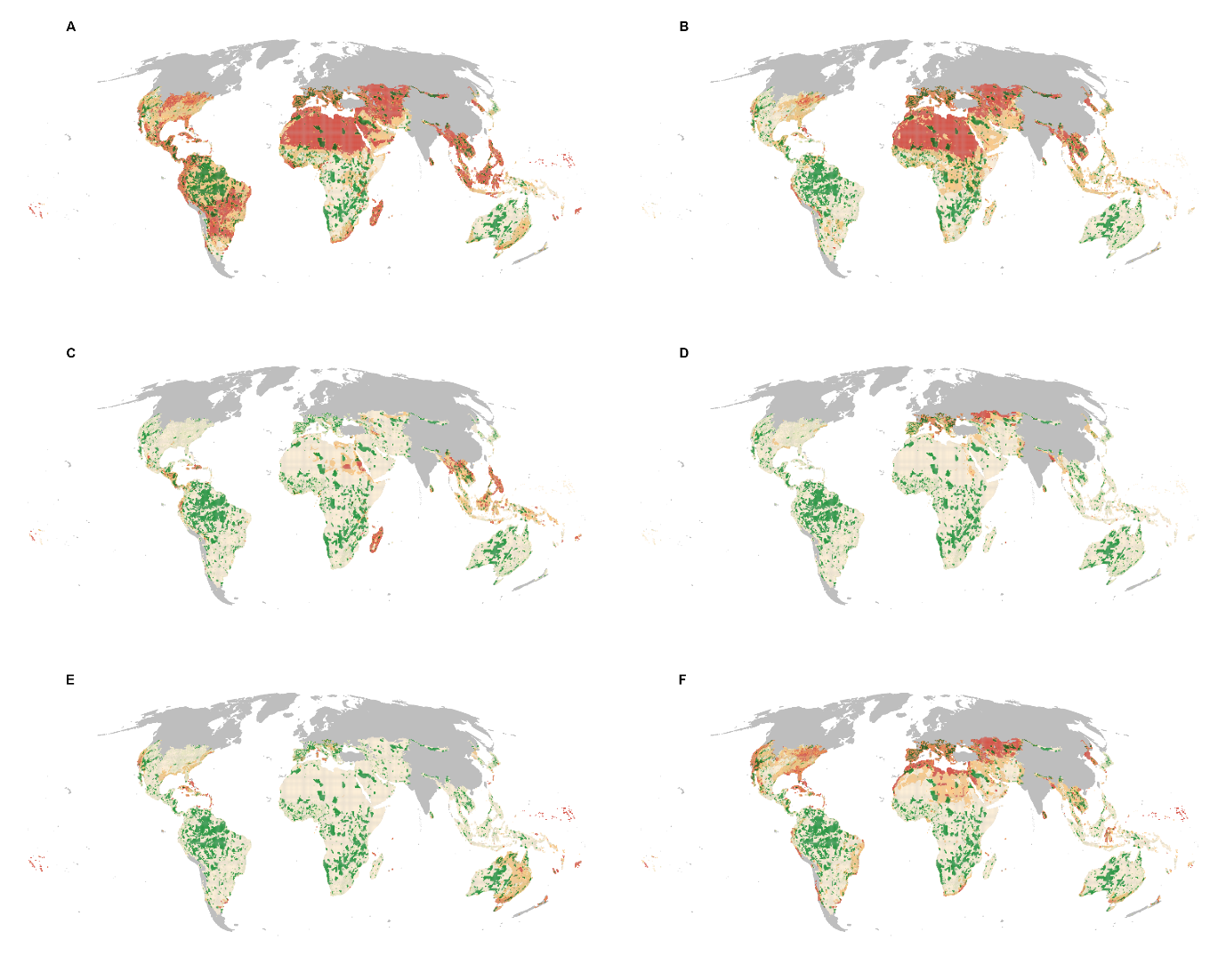

Figure S4. Bivariate maps for PA cover and threats to reptiles (A. Agriculture, B. Hunting, C. Logging, D. Pollution, E. Invasive species F. Urbanization)

**Increased Uncertainty Due to Range Sizes and Species Richness**

While the results offer valuable insights, it is crucial to acknowledge potential uncertainties. The median range sizes are largest for birds (average 26.3 km²) followed by mammals (7.1 km²). This raises uncertainty, particularly for birds, as the analysis considers average threats across large areas. Regions with the largest bird ranges, like Polynesia, Micronesia, Europe, Western Asia, Northern America, and Africa, should be interpreted cautiously due to this inherent uncertainty (see Supplementary Figures S5 & S6).

Furthermore, species richness can also influence the analysis. Areas with low species richness, like North America and Europe for amphibians and reptiles, or the High Arctic, North Africa, and Central Asia for birds and mammals, may require additional considerations. With fewer species present, threat data might not capture the full picture, potentially leading to higher uncertainty.

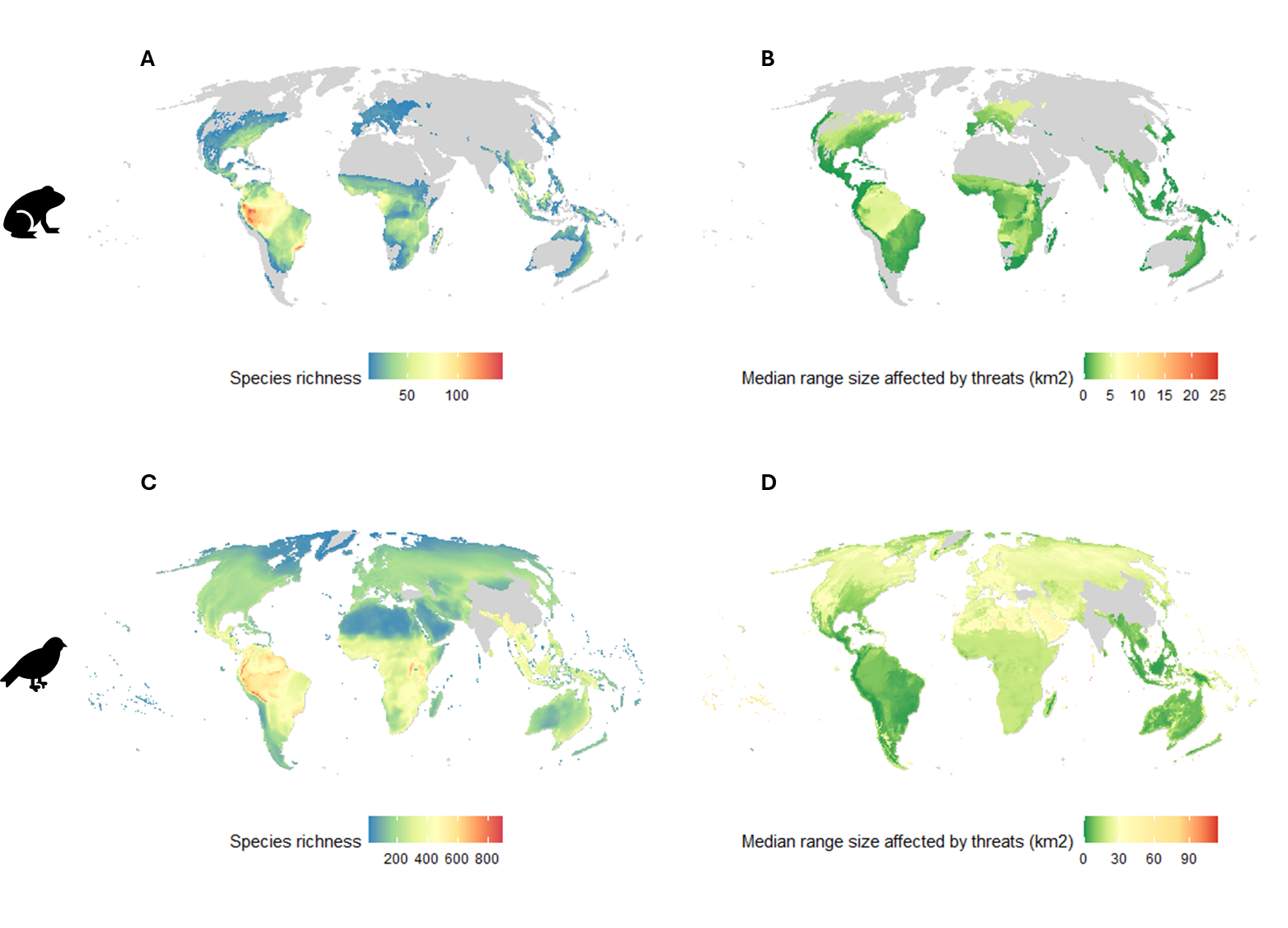

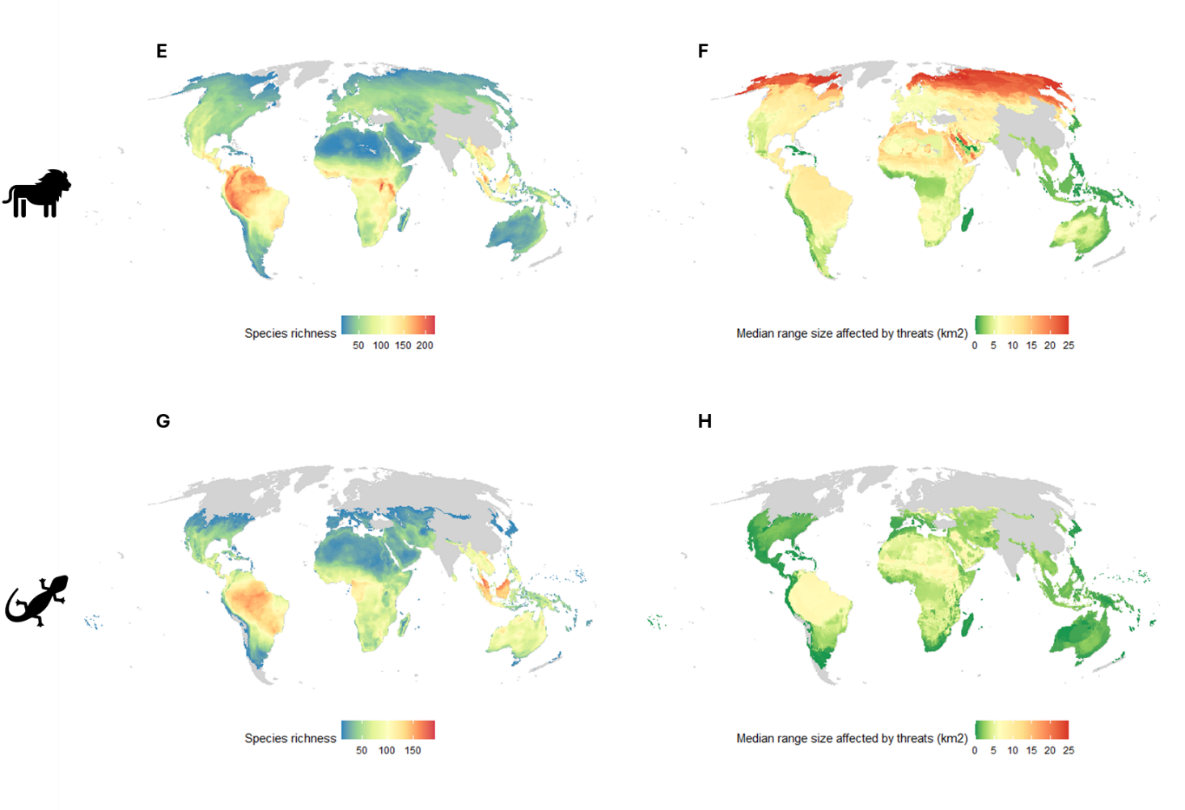

Figure S5. Species richness and median range sizes of threats for Amphibians (A -B), Birds (C – D), Mammals (E – F) and Reptiles (G – H)

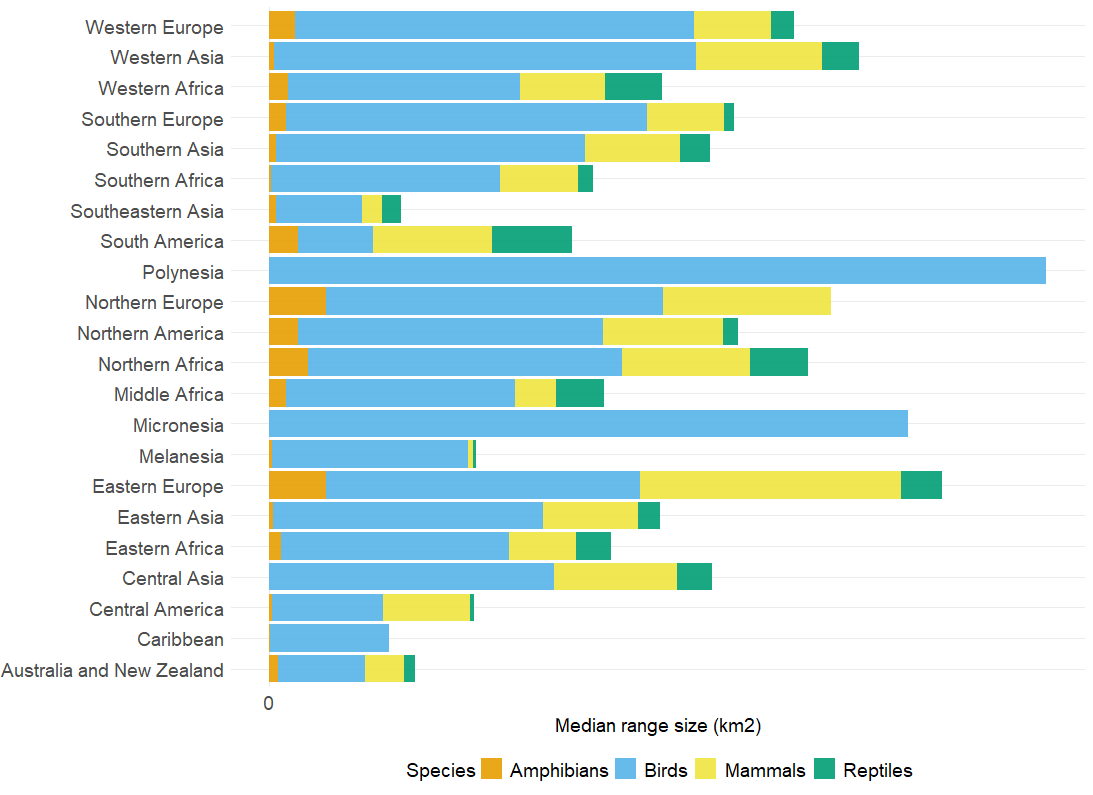

Figure S6. Regional median range size of species affected by threats
